# IFNG-producing self-reactive CD4^+^ T cells induce autoimmune adrenalitis in a mouse model of Addison’s disease

**DOI:** 10.1101/2025.08.11.669305

**Authors:** Arina Andreyeva, Juraj Michalik, Veronika Niederlova, Veronika Cimermanova, Ales Drobek, Radislav Sedlacek, Jan Prochazka, Juraj Labaj, Olha Fedosieieva, Waldemar Kanczkowski, Peter Draber, Andre Sulen, Ondrej Stepanek, Ales Neuwirth

**Affiliations:** Laboratory of Adaptive Immunity, Institute of Molecular Genetics of the Czech Academy of Sciences, Prague, Czechia; Faculty of Science, Charles University, Prague, Czechia; Czech Centre for Phenogenomics & Laboratory of Transgenic Models of Diseases, Institute of Molecular Genetics of the Czech Academy of Sciences, Vestec, Czechia; Department of Internal Medicine III, University Hospital Carl Gustav Carus, Technische Universität Dresden, Dresden, Germany; Department of Immunobiology, University of Lausanne, Epalinges, Switzerland; Department of Clinical Science, Faculty of Medicine, University of Bergen, Norway

**Keywords:** Addison’s disease, adrenals, autoimmunity, CYP11A1, CD4^+^ T cells, tolerance, IFNG, mouse model, AIRE

## Abstract

Autoimmune Addison’s disease (AD) is a rare but life-threatening disorder caused by immune-mediated destruction of the adrenal cortex, and progress in therapy has been limited by insufficient mechanistic insight. Here, we establish a model of Experimental Autoimmune Adrenalitis (EAA) that recapitulates key features of AD and reveals sex-dependent differences in disease manifestation within the model. Immunization with peptides derived from the adrenal self-antigen CYP11A1 induces corticosterone insufficiency. We show that autoimmune adrenalitis is driven by IFNG produced by self-reactive CD4⁺ T cells, promoting granulomatous inflammation in the adrenal cortex. Together, these findings identify IFNG as a central effector of autoimmune adrenalitis and suggest that targeting the IFNG pathway may represent a potential therapeutic strategy for AD.

**Significance:** Addison’s disease (AD) is a rare autoimmune disorder that destroys the adrenal cortex, yet its underlying mechanisms remain unknown. We developed a mouse model of Experimental Autoimmune Adrenalitis (EAA) that mirrors the hormonal and immunological features of AD. Our study reveals that IFNG-producing CD4⁺ T cells drive adrenal inflammation and dysfunction, identifying IFNG as a key pathogenic factor and a potential therapeutic target.

## Introduction

Addison’s disease (AD) is a primary adrenal insufficiency caused by autoimmune destruction of hormone-producing adrenocortical cells with a prevalence of 1 in 10,000 (1). AD leads to insufficient production of adrenocortical steroid hormones, which is fatal if untreated. Current management of AD consists of lifelong hormone replacement therapy (2). However, treatments do not fully mimic physiological hormone regulation, particularly under stress, leaving patients vulnerable to life-threatening adrenal crises (3). Moreover, hormone replacement therapy leads to adverse effects such as osteoporosis, increased blood pressure, obesity, or abnormal glucose metabolism (4, 5).

Historically, AD was frequently associated with tuberculosis, in which the mycobacterial spread to the adrenal glands caused direct granulomatous destruction of adrenal tissue (6). Currently, approximately 40% of AD cases are isolated autoimmune endocrinopathies, while the remaining cases occur in association with other autoimmune disorders, such as thyroiditis or type 1 diabetes (2, 7). Most notably, AD often manifests in patients with Autoimmune Polyendocrine Syndrome Type 1 (APS-1). This rare monogenic disorder is caused by mutations in the *AIRE* gene (8, 9), which leads to the emergence of autoreactive T cells in the periphery. Genetic predispositions to AD also involve polymorphisms in genes related to antigen presentation (HLA alleles DR3/DQ2 and DR4.4/DQ8; CIITA) and regulation of T-cell signaling (CTLA-4, PD-1, PTPN22) (10). Furthermore, emerging cancer immunotherapies, particularly immune checkpoint inhibitors, have been reported to induce AD as an adverse effect (11, 12).

The hallmark of AD is the presence of autoantibodies against the steroidogenic enzyme, 21-hydroxylase (21-OH, CYP21A2) (13, 14). These circulating antibodies serve as a predictive marker detectable even during the subclinical phase (15). Additionally, it is noteworthy that autoantibodies targeting cholesterol side-chain cleavage enzyme (CYP11A1) are less prevalent (16, 17) but observed in 70% of individuals with APS-1 (16, 18). Previous findings support an autoimmune-based mechanism in which self-reactive T cells play a crucial role in AD progression (19, 20). Both CD4^+^ and CD8^+^ T cells specific to immune-dominant epitopes of 21-OH are detected in the blood of the majority of patients with AD (20, 21). After in vitro antigen re-stimulation with 21-OH, a portion of CD4^+^ T cells produce IFNG suggesting that AD might be a T helper 1 cell (Th1)-associated autoimmune disorder (21). However, the mechanism underlying the destruction of adrenocortical cells remains elusive. Understanding of AD etiology has been hindered by the absence of a defined mouse model, and the unavailability of adrenal samples from AD patients, due to unjustified invasiveness of adrenal biopsies from living donors and the rarity of the disease.

Here, we present a mouse model of Experimental Autoimmune Adrenalitis that resembles AD. Using this model, we identified IFNG production by self-reactive CD4^+^ T cells as the key molecular mechanism driving the disease progression and, therefore, a potential therapeutic target for AD.

## Results

### Immunization with CYP11A1-derived peptides induces adrenal infiltration by heterogeneous CD4^+^ T cells

21-OH is the major self-antigen in human AD. However, no mouse models that induce an anti-21-OH immune response have been reported. Here, we sought an alternative self-antigen expressed both in adrenocortical cells and in the thymus in an AIRE-dependent manner. Our reanalysis of murine thymic single-cell RNA sequencing (scRNA-seq) data (22) revealed that *Cyp11a1* is expressed in medullary thymic epithelial cells (mTECs) in an AIRE-dependent manner, similar to the canonical tissue-restricted gene *Ins2*, whereas *Cyp21a1* (encoding mouse 21-OH) expression was nearly undetectable in the thymus (Figure 1A, Figure S1A). Based on these criteria, CYP11A1, previously identified as a target of autoantibodies in APS-1 patients (16, 18), emerged as a strong candidate autoantigen.

**Figure 1.**
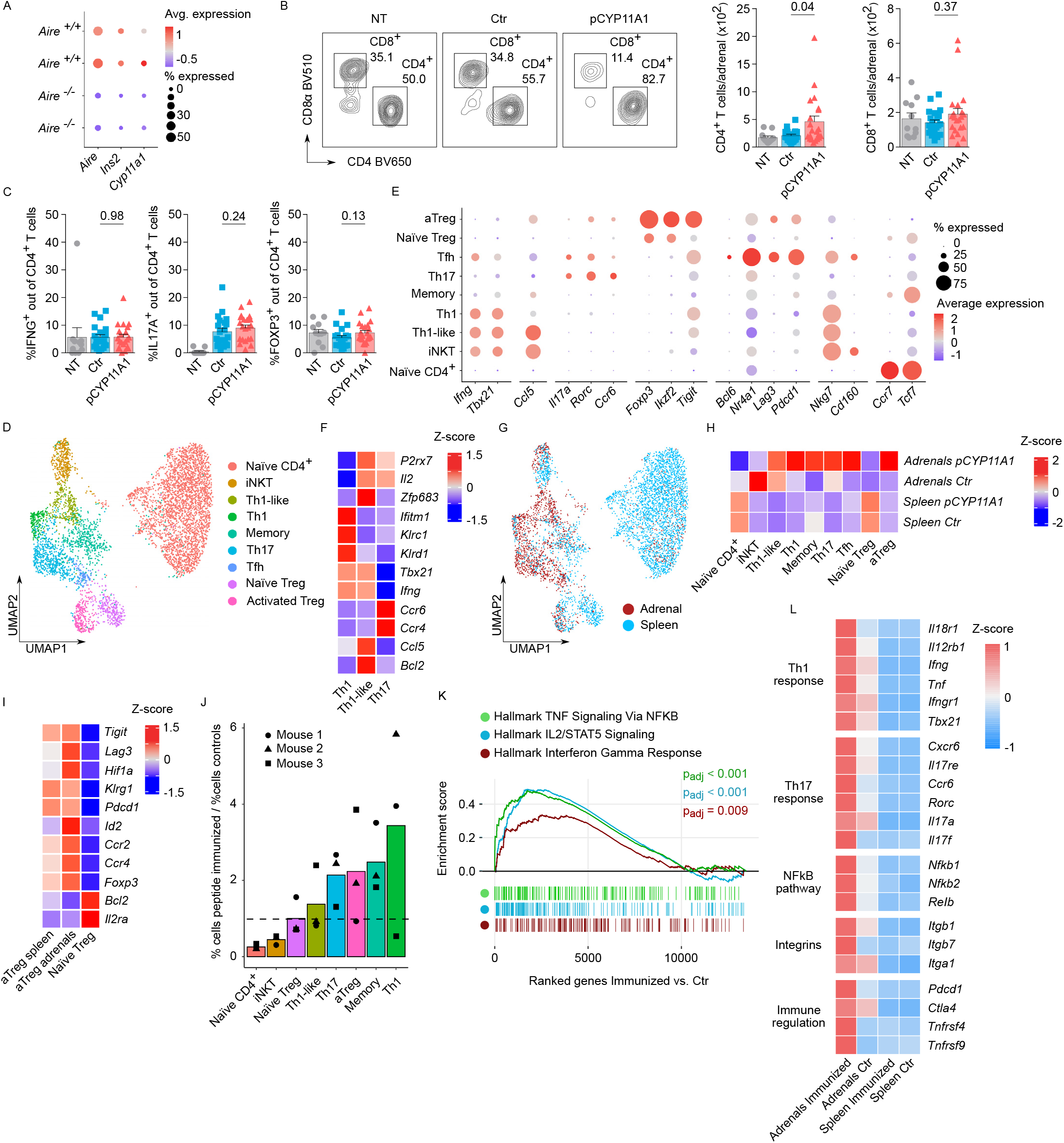
pCYP11A1 immunization recruits heterogeneous CD4⁺ T cells into the adrenals. **(A)** Expression of *Aire*, *Ins2*, and *Cyp11a1* in thymic epithelial cells (TECs) from *Aire^+/+^* and *Aire^−/−^* mice. Re-analysis of scRNA-seq data from 8-week-old female mice (22). Color indicates average expression; dot size indicates the proportion of expressing cells. **(B)** Representative gating and quantification of adrenal CD4⁺ and CD8⁺ T cells in non-treated (NT), control (Ctr), and pCYP11A1-immunized mice at day 14 p.i. (three independent experiments; n = 11-21 mice/ group). **(C)** Frequencies of IFNG⁺, IL17A⁺, and FOXP3⁺ adrenal CD4⁺ T cells after PMA/ionomycin re-stimulation (three independent experiments; n = 11-21 mice/group). (**B,C**) Statistical significance was determined using two-tailed Mann-Whitney test. Bars show mean ± SEM; symbols represent individual mice. **(D-L)** scRNA-seq analysis of CD4⁺ T cells from adrenal and spleen of pCYP11A1-immunized and control mice at day 14 p.i. Three mice exhibiting the highest adrenal CD4⁺ T-cell infiltration (based on flow cytometry; Figure S2A) were selected. Control adrenals were pooled from six CFA-immunized mice. **(D)** UMAP of CD4⁺ T-cell clusters from adrenal and spleen samples. **(E)** Dot plot of marker genes for cluster annotation. Data generated from adrenals. **(F)** Heatmap of Th1, Th1-like, and Th17 gene expression in immunized adrenals (row-normalized Z-scores). **(G)** UMAP showing tissue of origin. **(H)** Heatmap of CD4⁺ T-cell cluster abundance (column-normalized Z-scores). **(I)** Heatmap of naïve and activated Treg gene expression from spleen and adrenal samples (row-normalized Z-scores). **(J)** Ratio of CD4⁺ T-cell populations in peptide-immunized versus control mice. Dashed line indicates equal frequency. Data generated from adrenals. **(K)** GSEA of hallmark signatures in immunized versus control CD4⁺ T cells. P values were calculated by fgsea adaptive multilevel split Monte Carlo method with Benjamini-Hochberg correction. **(L)** Heatmap of selected genes in immunized and control spleen and adrenal (row-normalized Z-scores).

Using the Immune Epitope Database (23), we identified two CYP11A1 peptides (CYP11A1_150-164_ and CYP11A1_233-247_) predicted to bind I-A^b^ MHC class II molecule with the highest affinity (Figure S1B). B6 mice were immunized with both peptides (pCYP11A1) emulsified in complete Freund’s adjuvant (CFA). Two weeks after the immunization, we observed an increase in adrenal infiltration of CD4^+^ T cells in the immunized mice relative to CFA-only injected (Ctr) or non-treated (NT) controls (Figure 1B, Figure S1C). In contrast, adrenal CD8⁺ T-cell numbers remained comparable across all experimental groups (Figure 1B). On average, 8% of adrenal infiltrating CD4^+^ T cells expressed IL17A, 6% of them produced IFNG, and 6% were FOXP3^+^ regulatory T cells (Tregs) without differences between immunized and CFA-only control mice (Figure 1C, Figure S1D). Together, these data demonstrate that pCYP11A1 immunization induces adrenal infiltration by heterogeneous CD4⁺ T cells.

### Adrenal CD4^+^ T cells clonally expand and display diverse effector fates

To characterize adrenal infiltrating CD4^+^ T cells, we performed scRNA-seq combined with TCRαβ VDJ profiling of CD4^+^ T cells isolated from three mice with high CD4^+^ T-cell adrenal infiltration determined by flow cytometry at two weeks post-immunization (Figure S2A). As a control, adrenal CD4^+^ T cells from six CFA-immunized mice were pooled to obtain sufficient cell numbers (Figure S2A). Splenic CD4^+^ T cells from both groups of mice were analyzed as well.

Unsupervised clustering identified thirteen distinct clusters of adrenal and splenic CD4^+^ T cells (Figure S2B). Five clusters represented naïve CD4^+^ T cells and were merged into a single cluster for downstream analyses (Figure 1D). Among the *Cd44*^+^ antigen-experienced T cells (Figure S2C), eight clusters were annotated based on specific gene markers and/or VDJ usage: Th1 cells (*Ifng*^+^ *, Tbet*^+^), Th1-like cells (*Ifng*^+^, *Tbet*^+^, *Ccl5*^+^), Th17 cells (*Il17a*^+^*, Rorc*^+^*, Ccr6*^+^), activated Tregs (aTregs) (*Foxp3*^+^*, Ikzf2*^+^*, Tigit*^+^), naïve Tregs (*Foxp3*^+^*, Ikzf2*^+^), Tfh (*Bcl6*^+^*, Nr4a1*^+^*, Lag3*^+^*, Pdcd1*^+^), CD4^+^ iNKT cells (*Nkg7*^+^*, Cd160*^+^, CDR3α: CVVGDRGSALGRLHF) and memory T cells (*Ccr7*^+^, *Tcf7*^+^) (Figure 1E, Figures S2D-E). To our knowledge, the Th1-like IFNG-expressing cells, which differed from canonical Th1 cells by the expression of *Ccl5*, *P2rx7, Il2*, *Zfp683* (*Hobit*)*, Bcl2* and a lack of NK-related genes *Ifitm1*, *Klrc1*, and *Klrd1*, have not been previously described (Figures 1E-F, Figure S2D).

In contrast to the spleen with predominating naïve CD4^+^ T cells, the majority of adrenal CD4^+^ T cells displayed an effector phenotype (*Cd44*^+^*, Icos*^+^*, Sell*⁻*, Ccr7*⁻) (Figures 1G-H, Figure S2C). In the control adrenals, CD4^+^ T cells were predominantly composed of iNKT cells with a limited presence of effector cells (Figure 1H).

The naïve Treg cluster detected in the spleen was almost absent in the adrenals (Figures 1G-H). Instead, adrenal Tregs exhibited an activated phenotype, as demonstrated by the changes in gene expression, including increased expression of *Tigit*, *Lag3*, *Hif1a*, *Klrg1*, *Pdcd1*, *Id2, Ccr2*, *Ccr4*, *Foxp3*, and lower expression of *Bcl2* and *Il2ra* (Figures 1E, I).

Th17 and memory CD4^+^ T cells were enriched in all three immunized mice in comparison to the control sample, which comprised pooled adrenal CD4^+^ T cells from CFA-only treated mice (Figure 1J, Figure S2F). Inversely, the frequencies of naïve CD4^+^ cells and iNKT cells were higher in the control samples than in the immunized mice. Two mice showed strongly enriched Th1 cells, whereas the third one had elevated Th1-like and aTreg cells illustrating the mouse-to-mouse variability (Figure 1J).

To complement phenotypic observations with gene-level analysis, we performed gene set enrichment analysis (GSEA) on adrenal CD4⁺ T cells from peptide-immunized versus control mice. This revealed enrichment of gene signatures associated with inflammatory cytokines TNF, IL-2, and IFNG after pCYP11A1 immunization (Figure 1K). Additionally, differential expression analysis across all experimental groups showed upregulation of proinflammatory Th1- and Th17-related genes as well as genes associated with T-cell activation (Figure 1L). These transcriptional changes are consistent with the increased abundance of Th1 and Th17 effector subsets identified by scRNA-seq.

In the next step, we identified clonally expanded families based on their shared TCRα and TCRβ sequences (Figure 2A). The clonally expanded T cells were present primarily within effector populations in the adrenal samples from all three pCYP11A1-immunized mice and only marginally in the spleen (Figure 2B, Figure S2G). Th17, Th1-like, Tfh, Th1, aTreg and memory cells were the subsets with substantial clonal expansion (Figure 2C).

**Figure 2.**
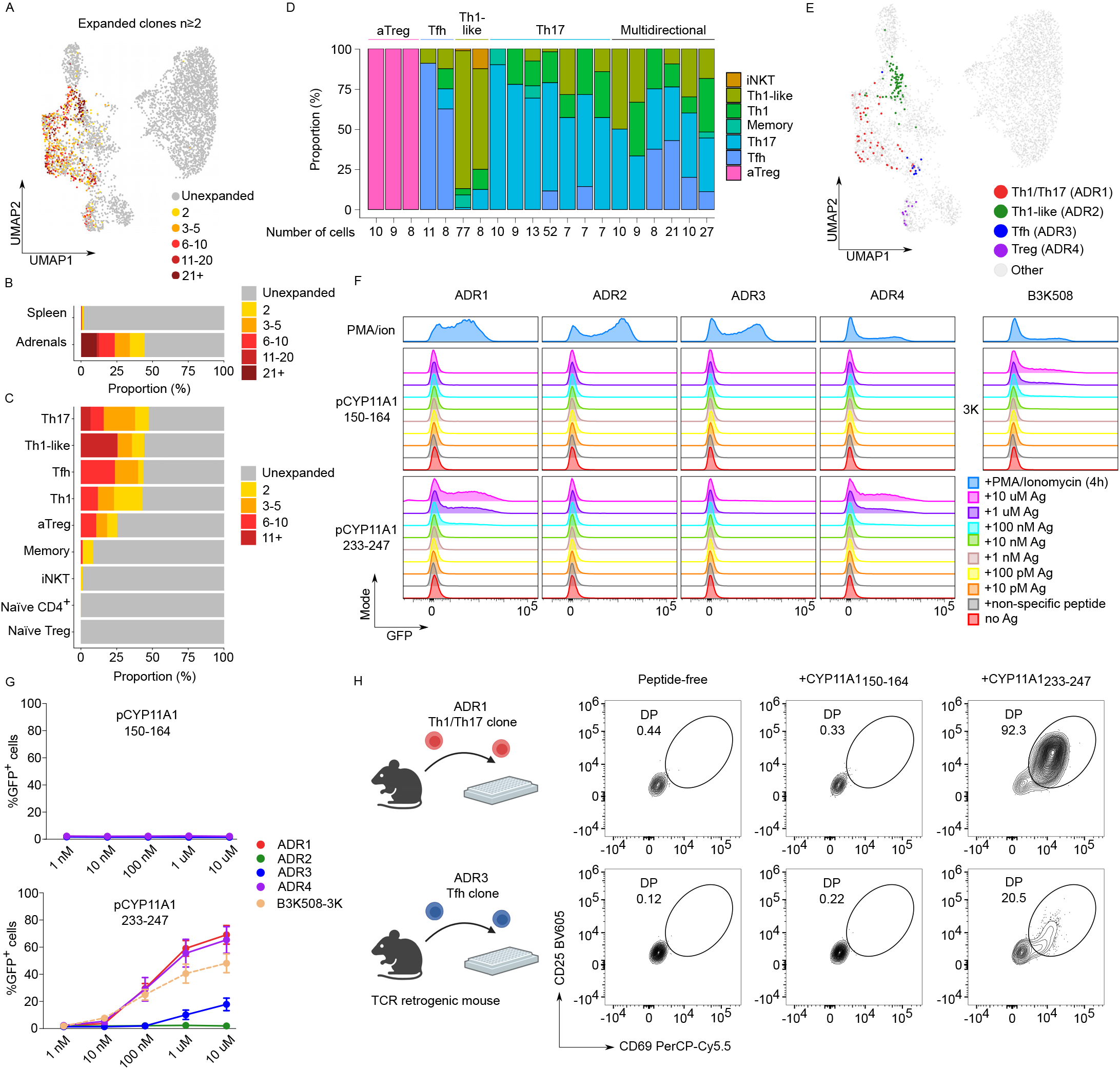
Adrenal CD4⁺ T cells are clonally expanded and include CYP11A1-reactive clones. **(A)** UMAP of CD4⁺ T cells from adrenal glands and spleens of pCYP11A1-immunized mice showing expanded T-cell clones (≥2 cells sharing the same TCRαβ). **(B)** Distribution of expanded CD4⁺ T-cell clones between adrenal glands and spleens. **(C)** Distribution of expanded CD4⁺ T-cell clones across defined T-cell subsets. Clone size was calculated separately for each subset. **(A-C)** Color indicates clone size (number of cells sharing the same TCRαβ). **(D)** Composition of the 20 largest expanded CD4⁺ T-cell clones. Clones are arranged by effector phenotype diversity, from polarized (left) to multidirectional (right); cell numbers are indicated on the x-axis. Colors indicate effector phenotype; dominant phenotypes (≥50%) are shown above. **(E)** UMAP highlighting four expanded CD4⁺ T-cell clones (ADR1-ADR4) among effector populations. Colors indicate clone identity. **(F)** Representative histograms of NFAT-GFP reporter activation in A5 hybridomas expressing ADR1-ADR4 TCRs stimulated with serial dilutions of CYP11A1_150-164_ or CYP11A1_233-247_. B3K508 A5 hybridomas stimulated with 3K peptide served as a positive control, whereas peptide-free cultures and irrelevant peptides at the highest concentration served as negative controls. PMA/ionomycin served as a positive stimulation control. Colors indicate peptide concentration. Representative of three independent experiments. **(G)** Quantification of GFP⁺ cells corresponding to (F). Data are mean ± SEM from three independent experiments. Colors indicate individual T-cell clones. **(H)** Experimental design for testing antigen specificity of two retrogenic CD4⁺ T-cell clones. CD69 and CD25 expression was measured after stimulation with CYP11A1_150-164_ or CYP11A1_233-247_; peptide-free cultures served as controls. Representative of three independent experiments. Created with BioRender.com

Most clones exhibited a bias toward a specific lineage phenotype, with a predominance of the Th17 lineage, followed by aTreg, Tfh, and Th1-like subsets. Approximately one-third of the clones displayed a multidirectional effector phenotype. Notably, many clones with a dominant Th17 bias also gave rise to Th1 cells (Figure 2D).

To investigate antigen specificity, four representative clones with distinct effector profiles were selected (Figure 2E). Their TCRs were expressed in an NFAT-GFP reporter T-cell hybridoma line (A5) (24, 25), and reactivity to pCYP11A1 was assessed using an antigen presentation assay (Figures 2F-G, Figure S2H). Three clones: ADR1 (Th1/Th17), ADR3 (Tfh), and ADR4 (Treg)-exhibited specificity for the CYP11A1_233-247_ epitope, with differing levels of activation, whereas the ADR2 (Th1-like) clone did not respond to either of the pCYP11A1 peptides tested (Figures 2F-G). Notably, ADR1 and ADR4 clones displayed strong antigen reactivity comparable to hybridoma cells expressing a control B3K508 TCR in response to its cognate exogenous 3K peptide (26, 27) (Figure 2G).

In addition, we generated retrogenic mice expressing individual TCRs. Two clones, ADR1 and ADR3, successfully gave rise to monoclonal CD4⁺ T-cell populations in vivo. In subsequent antigen presentation assays, both ADR1 and ADR3-expressing T cells specifically responded to antigen-presenting cells loaded with CYP11A1_233-247_ peptide, but not to CYP11A1_150-164_ or unloaded controls (Figure 2H) confirming antigen specificity. Notably, ADR3 exhibited lower reactivity compared to ADR1, consistent with our previous observations in A5 reporter cells.

These results demonstrate that pCYP11A1 immunization induces clonal expansion of phenotypically diverse autoreactive CD4^+^ T cells in the adrenal gland.

### The development of the Experimental Autoimmune Adrenalitis model

Because adrenal CD4^+^ T-cell infiltration was variable after a single immunization, we introduced a booster injection containing pCYP11A1, poly(I:C), and anti-CD40 antibodies (28). One week later, flow cytometry revealed enhanced adrenal infiltration by CD4^+^ T cells together with CD8^+^ T cells, B cells, and CD11b^+^ myeloid cells (Figure 3A, Figure S3A). The magnitude of CD4⁺ T-cell infiltration was comparable between male and female mice (Figure 3B). Because this protocol was more robust and reproducible, it was used in all subsequent experiments and is referred to hereafter as Experimental Autoimmune Adrenalitis (EAA).

**Figure 3.**
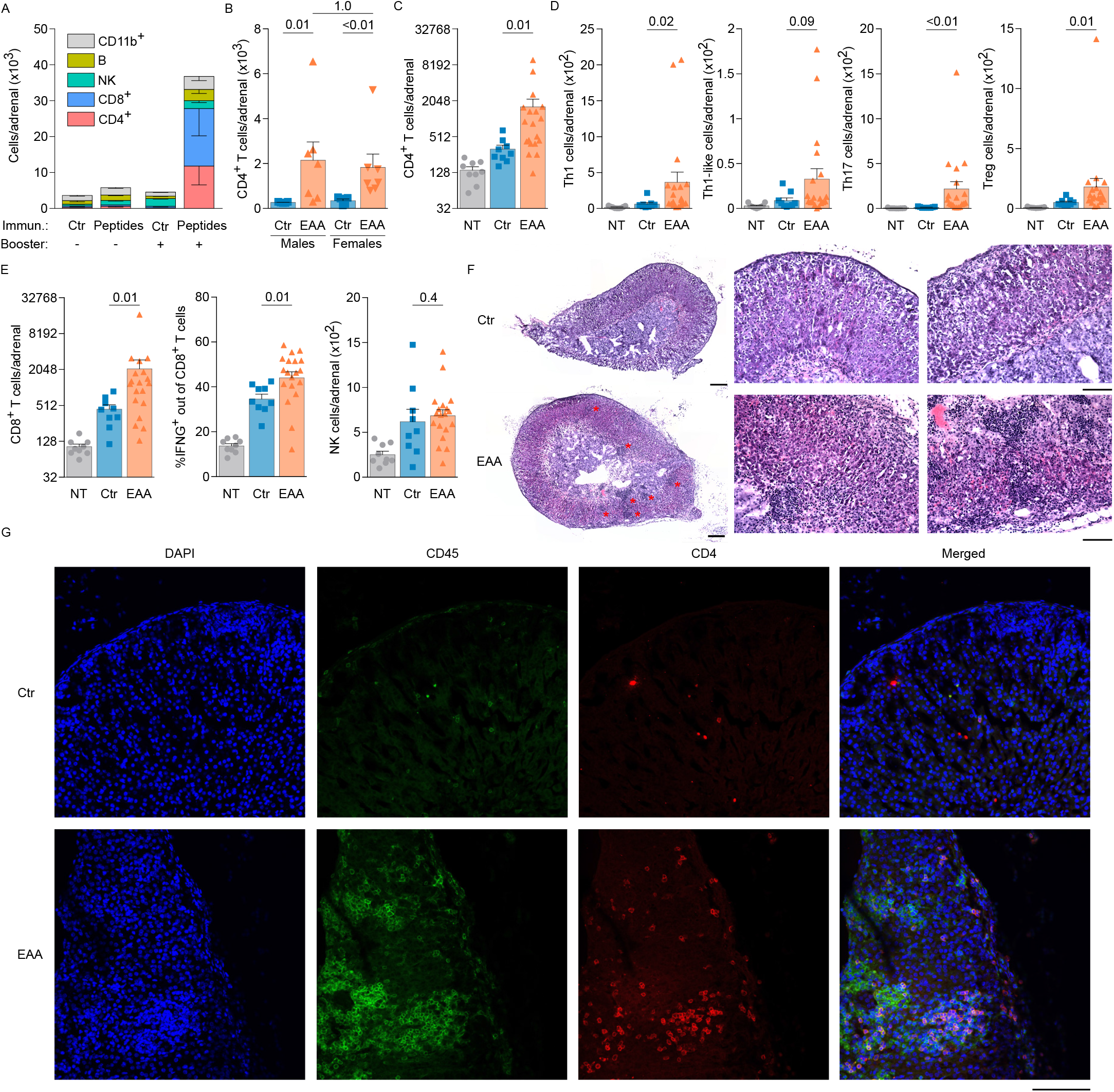
Experimental autoimmune adrenalitis (EAA) model. **(A)** Quantification of total immune cells in adrenal glands of control and pCYP11A1-immunized mice (± booster) at day 25 p.i. (n = 5-6 mice/group). Bars show mean ± SEM; colors indicate immune cell subsets. **(B)** Sex-dependent quantification of adrenal CD4⁺ T cells at 3-4 weeks p.i. (n = 5-7 mice/group). Groups: control (Ctr) and EAA. **(C,D)** Quantification of adrenal CD4⁺ T cells and effector CD4⁺ T-cell subsets at 3-4 weeks p.i. **(E)** Quantification of adrenal CD8⁺ T cells, IFNG⁺ CD8⁺ T cells, and NK cells. **(C-E)** Statistical significance was determined using the two-tailed Mann-Whitney test; P values are indicated. Bars show mean ± SEM; symbols represent individual mice (two independent experiments; n = 9-19 mice/group). Groups: non-treated (NT), control (Ctr), and EAA. **(F)** Representative H&E-stained adrenal sections at 3 weeks p.i. (one experiment; n = 4-5 mice/group). Whole-section images (scale bars, 200 μm) and magnified views (scale bar, 100 μm). Asterisks indicate leukocyte infiltrates. **(G)** Representative immunofluorescence images of adrenal sections stained for DAPI (nuclei, blue), CD45 (green), and CD4 (red) (one experiment). Scale bar, 100 μm.

Using flow cytometry informed by the scRNA-seq analysis (Figures 1D-E, Figure S2D), we identified Th1 (IFNG^+^ CCL5⁻), Th1-like (IFNG^+^ CCL5^+^), Th17 (IL17A^+^), and Treg (FOXP3^+^) populations among adrenal CD4⁺ T cells after PMA/ionomycin restimulation (Figure S3B). EAA mice exhibited increased numbers of adrenal CD4⁺ T cells and all analyzed effector subsets compared with controls (Figures 3C-D, Figure S3C). Among these populations, only Th17 cells showed an increased relative frequency (Figure S3C). IFNG^+^ CD8^+^ T cells, but not NK cells, were significantly increased in EAA mice; whereas mock immunization induced a weaker CD8^+^ T-cell response (Figure 3E).

Histological analysis showed that leukocytes localized predominantly to the zona fasciculata, with extension toward the juxtamedullary region and medulla (Figure 3F). CD45^+^ and CD4^+^ immunofluorescence confirmed the presence of infiltrating leukocytes and T cells within the adrenal cortex (Figure 3G). This distribution resembles the pattern of adrenal infiltration reported in patients with AD (29, 30).

Overall, our data demonstrated that the EAA protocol elicits a strong immune response, characterized by specific recruitment of inflammatory cells to the adrenal cortex, which is the site of extensive steroidogenesis.

### CYP11A1-driven autoimmunity induces adrenal pathology and dysfunction

We investigated whether immune cell infiltration of the adrenal gland affects its morphology and endocrine function. Because adrenal morphology and corticosterone production are sexually dimorphic (31), initial experiments were performed in male mice. Three to five months after EAA induction, H&E staining revealed enlarged multinucleated foamy cells (Figure 4A) with high lipid content (Figure 4B) and green autofluorescence (Figure 4C). These lipid-rich structures lacked CYP11A1 expression (Figure 4C). They were F4/80^+^, indicating a macrophage component (Figure 4D). These F4/80^+^ multinucleated lesions morphologically resembled the granulomas described in tuberculosis-associated adrenalitis (32), although their precise relationship to infectious granulomas remains unclear. Granulomas occupied approximately 2% of the adrenal cortex at 3 months and 5% at 5 months (Figure 4E).

**Figure 4.**
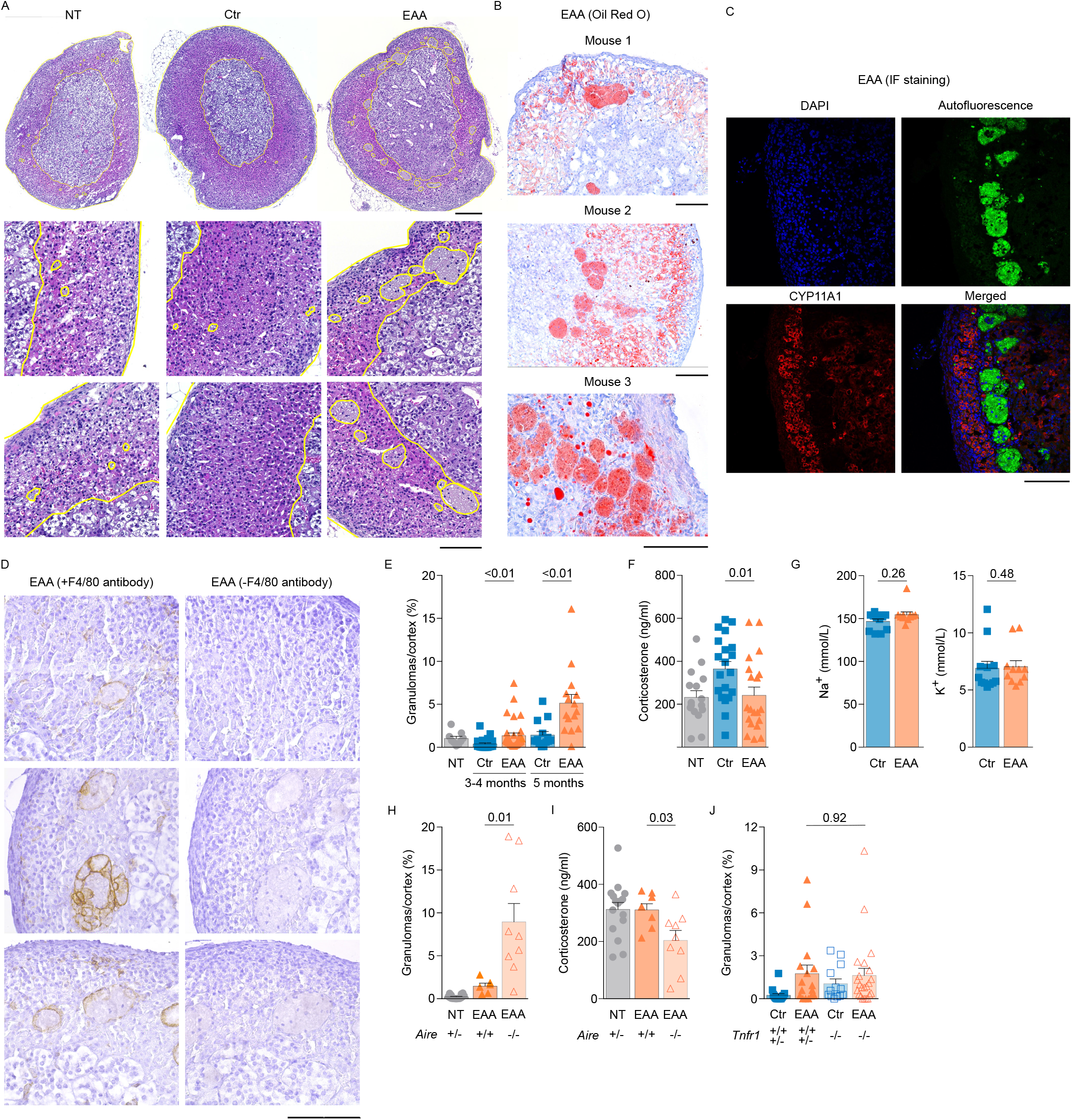
Long-term adrenal pathology in EAA. **(A)** Representative H&E-stained adrenal sections 3-5 months p.i. (n = 9 NT, 33-47 Ctr and EAA mice). Whole sections (scale bar, 200 μm) and higher-magnification views (scale bar, 100 μm). Adrenal borders, medulla, and multinucleated foamy cells are highlighted in yellow. **(B)** Representative Oil Red O staining of adrenal sections from EAA mice (one experiment; n = 6). Scale bar, 100 μm. **(C)** Representative immunofluorescence of adrenal sections from EAA mice stained for DAPI (nuclei, blue) and CYP11A1 (red) (one experiment; n = 6). Multinucleated foamy cells appear green due to lipid autofluorescence. Scale bar, 100 μm. **(D)** Representative F4/80 staining showing macrophage infiltration in adrenal sections from EAA mice; the right panel shows the no-primary-antibody control (one experiment; n = 8). Middle and bottom panels show different fields of the same section. Scale bar, 100 μm. **(E)** Cortical granuloma frequency in B6 mice 3-4 months p.i. (four independent experiments; n = 24-32 mice/group) and 5 months p.i. (three independent experiments; n = 9-15 mice/group). **(F)** Serum corticosterone after ACTH stimulation in B6 mice 5 months p.i. (three independent experiments; n = 16-20 mice/group). **(G)** Basal serum sodium and potassium in male B6 mice 5 months p.i. (two independent experiments; n = 11-12 mice/group). **(H,I)** Cortical granuloma frequency and serum corticosterone after ACTH stimulation in *Aire*^+/+^, *Aire*^−/−^, and NT (*Aire*^+/−^) mice 5 months p.i. (two independent experiments; n = 6-16 mice/group). **(J)** Cortical granuloma frequency in *Tnfr1*^+/+^, *Tnfr1*^+/−^, and *Tnfr1*^−/−^ mice 5 months p.i. (five independent experiments; n = 13-23 mice/group). **(E-J)** Statistical significance was determined using the two-tailed Mann-Whitney test; P values are indicated. Bars show mean ± SEM; symbols represent individual mice.

In the next step, we examined the adrenal function in the EAA mice. We measured blood corticosterone levels following ACTH stimulation, which is the diagnostic test for adrenal sufficiency routinely used in human patients (33). EAA male mice exhibited a significant reduction in corticosterone production compared to peptide-free immunized control mice (Figure 4F). Normal blood sodium and potassium levels, along with an intact zona glomerulosa, indicated preserved mineralocorticoid production (Figure 4G). Together, these findings demonstrate that EAA induces granulomatous adrenal inflammation and corticosterone insufficiency in males.

### AIRE deficiency exacerbates adrenal pathology in the EAA model

Since AIRE-dependent self-antigen expression in the thymus contributes to T-cell tolerance, and CYP11A1 is absent in AIRE-deficient mTECs (22), we investigated whether *Aire*^-/-^ mice would exhibit increased susceptibility to adrenal pathology in our model. Indeed, both hallmarks of EAA - granuloma formation and reduced corticosterone production - were significantly enhanced in the *Aire*^-/-^ mice (Figures 4H-I). These findings underscore the critical role of AIRE-mediated central tolerance in protecting against adrenal autoimmunity, highlighting that the absence of thymic expression of CYP11A1 increases susceptibility to autoimmune adrenal inflammation.

Next, we tested whether tumor necrosis factor (TNF), a pro-inflammatory cytokine contributing to several autoimmune and inflammatory disorders (34), is involved in the EAA induction. Although TNF signaling appeared relevant in our transcriptional analyses, *Tnfr1*^-/-^ mice lacking the major TNF receptor displayed similar levels of granuloma formation as TNFR-sufficient mice, suggesting that the TNF signaling is not essential for EAA induction (Figure 4J).

### CYP11A1-restimulated CD4⁺ T cells accelerate adrenal pathology in adoptive transfer EAA

To determine whether autoreactive CD4⁺ T cells are sufficient to induce disease, we adoptively transferred pCYP11A1-restimulated CD44⁺ CD4⁺ T cells into *Cd3e*^-/-^ or *Rag2*^-/-^ male recipients before EAA induction (Figure 5A). In both recipient strains, transferred CD4⁺ T cells accelerated disease, resulting in impaired corticosterone production and extensive granuloma formation three months after EAA induction (Figures 5B-D). Histology and flow cytometry confirmed increased leukocyte infiltration of the zona fasciculata together with increased adrenal CD4⁺ T-cell numbers (Figures 5D-E). Corticosterone production showed only a mild inverse correlation with granuloma burden and CD4⁺ T-cell infiltration (Figure S4A).

**Figure 5.**
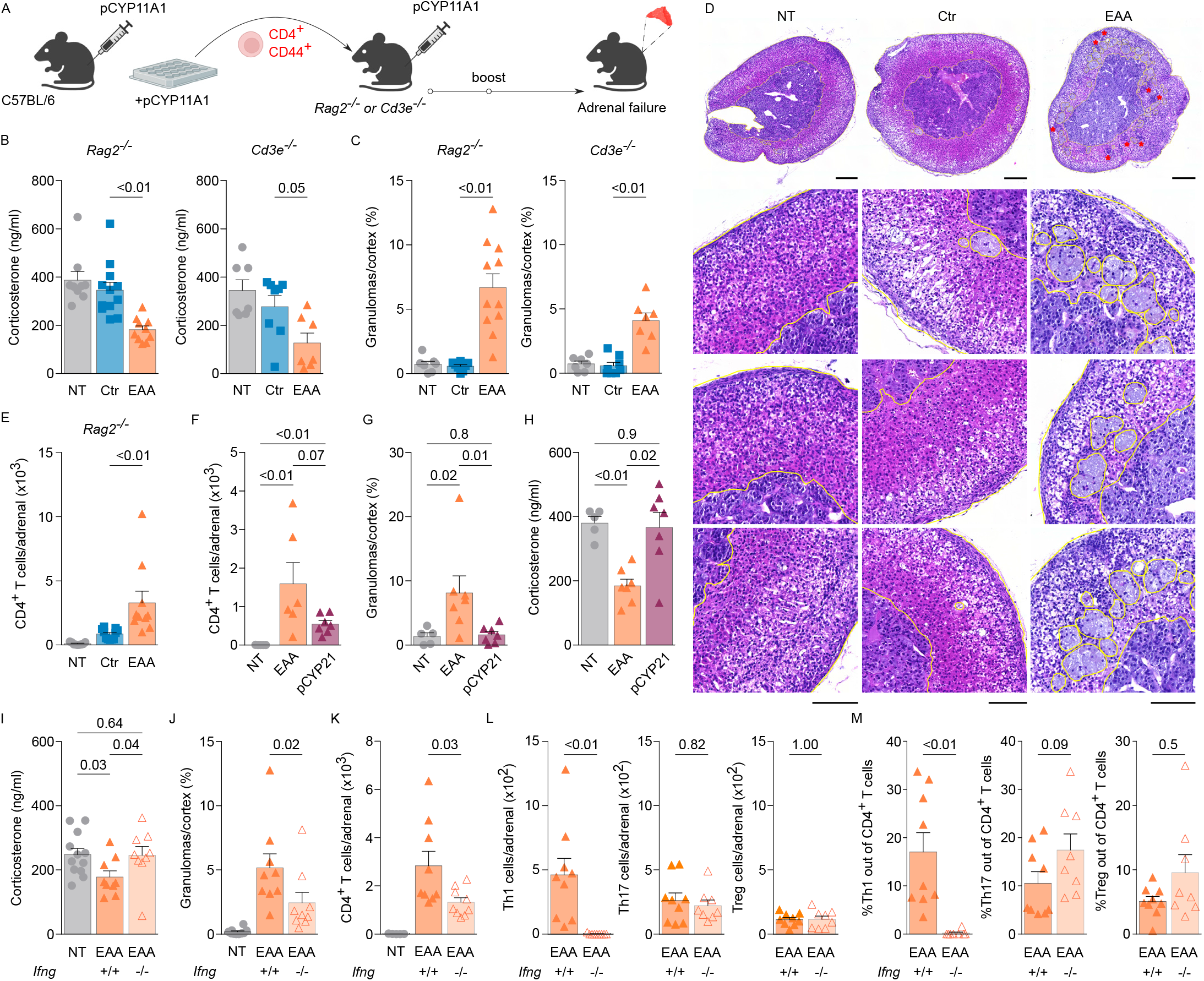
CD4⁺ T cell-derived IFNG drives pathology in the adoptive EAA model. **(A)** Adoptive EAA model design. CYP11A1-restimulated CD4⁺ T cells were transferred into *Rag2*^−/−^ or *Cd3e*^−/−^ recipients. Created with BioRender.com. **(B)** Serum corticosterone after ACTH stimulation in *Rag2*^−/−^ (two independent experiments; n = 9-12 mice/group) and *Cd3e*^−/−^ (two independent experiments; n = 7-8 mice/group) recipients 3 months post-transfer. **(C)** Cortical granuloma frequency in *Rag2*^−/−^ (two independent experiments; n = 7-11 mice/group) and *Cd3e*^−/−^ (two independent experiments; n = 7-8 mice/group) recipients 3 months post-transfer. **(D)** Representative H&E-stained adrenal sections from *Rag2*^−/−^ recipients (two independent experiments; n = 7-11 mice/group). Whole sections (scale bar, 200 μm) and magnified views (scale bar, 100 μm). Adrenal borders, medulla, and granulomas are highlighted in yellow; asterisks indicate infiltrating leukocytes. **(E)** Quantification of adrenal CD4⁺ T-cell infiltration in *Rag2*^−/−^ recipients 3 months post-transfer (two independent experiments; n = 10-11 mice/group). **(F-H)** Comparison of CYP11A1- and CYP21-specific immunization. CD4⁺ T cells from pCYP11A1- or pCYP21-immunized B6 mice were restimulated with corresponding peptides and transferred into *Cd3e*^−/−^ recipients (two independent experiments; n = 5-7 mice/group; analysis at 3 months post-transfer). **(F)** Quantification of CD4⁺ T-cell infiltration. **(G,H)** Cortical granuloma frequency and serum corticosterone after ACTH stimulation. *Cd3e*^−/−^ mice without transfer served as NT controls. **(I-M)** IFNG requirement in adoptive EAA. CD4⁺ T cells from *Ifng*^+/+^ or *Ifng*^−/−^ donors were transferred into *Rag2*^−/−^ recipients (two independent experiments; n = 9-12 mice/group; analysis at 3 months post-transfer). **(I,J)** Serum corticosterone after ACTH stimulation and cortical granuloma frequency. **(K-M)** CD4⁺ T-cell infiltration and effector subset distribution. NT mice were not analyzed in K-M due to absence of endogenous T cells. **(B,C,E-M)** Statistical significance was determined using two-tailed Mann-Whitney test; P values are indicated. Bars show mean ± SEM; symbols represent mice.

Since CYP21 is the dominant self-antigen in AD, we investigated whether CYP21-derived peptides (pCYP21) could induce autoimmune adrenalitis in the adoptive transfer setup. Initial experiments using two CYP21-derived peptides (CYP21_143-158_, and CYP21_344-359_) identified CYP21_344-359_ as the immunogenic epitope in the peripheral response (Figures S4B-C). A third peptide (CYP21_372-386_) predicted to have the highest binding affinity for the I-A^b^ MHC class II molecule, was subsequently designed using the Immune Epitope Database and used together with CYP21_344-359_ in adoptive transfer experiments. Despite its ability to induce a strong peripheral IFNG response, pCYP21 elicited weaker CD4^+^ T cell adrenal infiltration than pCYP11A1 (Figure 5F) and failed to induce granuloma formation or corticosterone insufficiency (Figures 5G-H). Absolute numbers and frequencies of Th1, Th1-like, Th17, and Treg populations were not significantly altered following pCYP21-mediated transfer (Figure S4D). In both conditions, IFNG-producing CD4⁺ T cells predominated, accounting for approximately 30-40% of total CD4⁺ T cells (Figure S4D). These findings demonstrate that pCYP11A1, but not pCYP21, acts as a pathogenic self-antigen in the EAA model, promoting adrenal infiltration, granuloma formation, and corticosterone insufficiency.

Together, these findings demonstrate that autoreactive CD4⁺ T cells are sufficient to induce EAA, whereas CD8⁺ T cells and B cells are dispensable.

### CD4⁺ T cell-derived IFNG is required for autoimmune adrenal insufficiency

To determine the role of CD4⁺ T cell-derived IFNG, we adoptively transferred CYP11A1-restimulated *Ifng*^-/-^ CD4^+^ T cells. Unlike WT CD4^+^ T cells, *Ifng*^-/-^ CD4⁺ T cells failed to induce adrenal insufficiency, as recipient mice maintained normal ACTH-stimulated corticosterone production and developed fewer adrenal granulomas (Figures 5I-J). Flow cytometric analysis of adrenal CD4^+^ T cells revealed increased cellularity and a dominant Th1 phenotype following transfer of WT, but not *Ifng*^-/-^ CD4^+^ T cells (Figures 5K-M). Together, these findings identify CD4^+^ T cell-derived IFNG as a critical mediator of autoimmune adrenal insufficiency.

### Sex-dependent uncoupling of adrenal pathology and endocrine dysfunction

We examined whether the EAA model could be established in females. Following two-step immunization, females neither developed adrenal granulomas nor showed impaired corticosterone production (Figures 6A-C).

**Figure 6.**
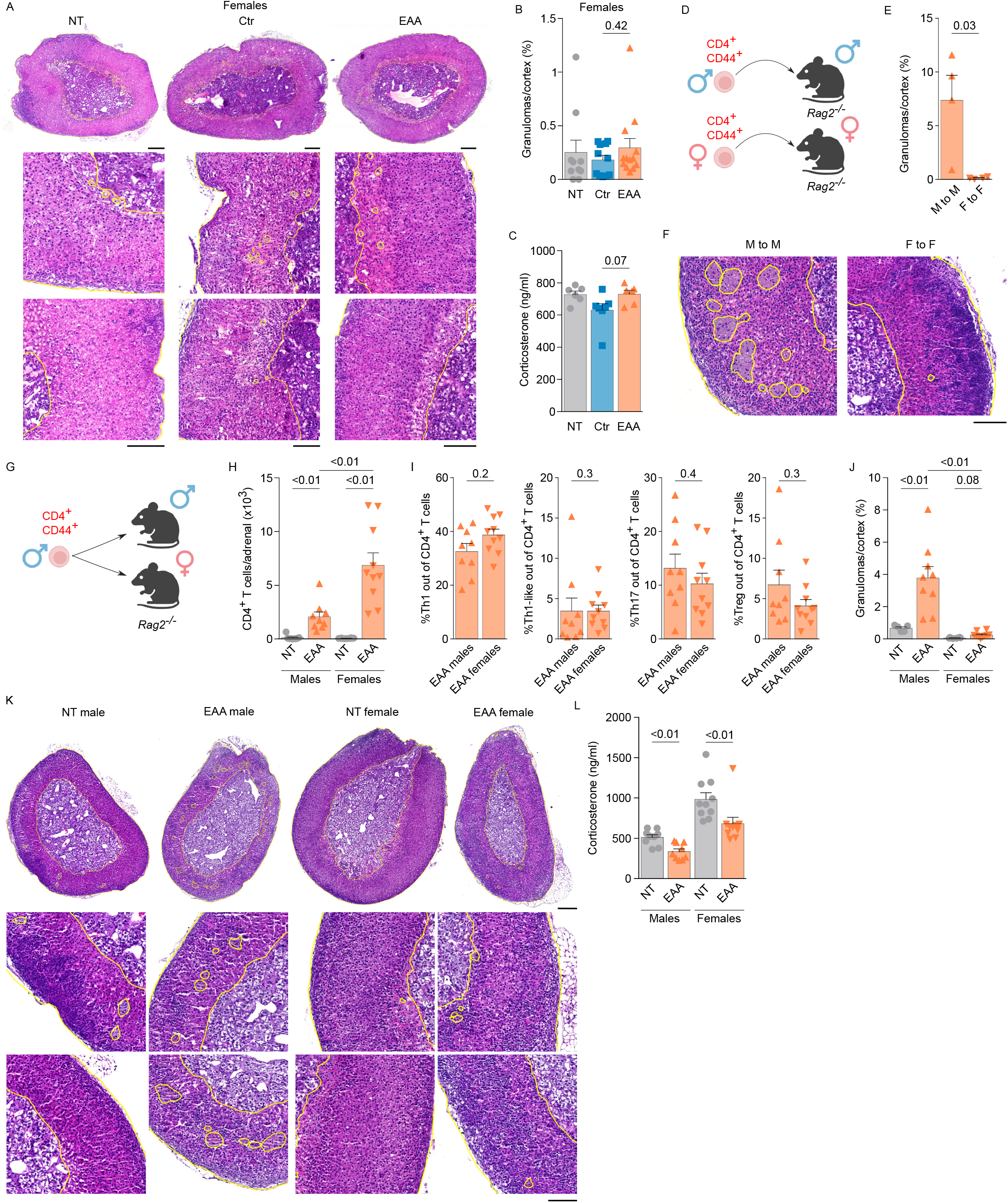
Sex-specific differences in EAA severity with male-restricted granuloma formation and limited functional impairment in females. **(A)** Representative H&E-stained adrenal sections from female B6 mice 5 months p.i. (two independent experiments; n = 10-13 mice/group). Whole-organ sections (scale bar, 200 μm) and magnified views (scale bar, 100 μm). Adrenal borders, medulla, and granulomas are highlighted in yellow. **(B)** Cortical granuloma frequency in female B6 mice 5 months p.i. (two independent experiments, n = 10-13 mice/group). **(C)** Serum corticosterone after ACTH stimulation in female B6 mice 5 months p.i. (n = 6-7 mice/group). **(D)** Experimental design of the adoptive EAA model. CYP11A1-restimulated CD4⁺ T cells from male or female donors were transferred into sex-matched *Rag2*^−/−^ recipients. Created with BioRender.com. **(E,F)** Sex-matched adoptive transfer model; analysis at 3 months post-transfer. **(E)** Cortical granuloma frequency in male and female *Rag2*^−/−^ recipients (n = 4 mice/group). **(F)** Representative H&E-stained adrenal sections (scale bar, 100 μm); adrenal borders, medulla, and granulomas are highlighted in yellow. **(G)** Experimental design of the adoptive EAA model using male donor CD4⁺ T cells transferred into male or female *Rag2*^−/−^ recipients. Created with BioRender.com. **(H-L)** Analysis of male donor CD4⁺ T-cell transfer into male and female *Rag2*^−/−^ recipients 3 months post-transfer (two independent experiments; n = 9-10 mice/group). **(H-I)** Quantification of CD4⁺ T-cell infiltration and effector CD4⁺ T-cell subsets. **(J)** Cortical granuloma frequency. **(K)** Representative H&E-stained adrenal sections. Whole-organ sections (scale bar, 200 μm) and magnified views (scale bar, 100 μm); adrenal borders, medulla, and granulomas are highlighted in yellow. **(L)** Serum corticosterone after ACTH stimulation. **(B,C,E,H-J,L)** Statistical significance was determined using two-tailed Mann-Whitney test; P values are indicated. Bars show mean ± SEM; symbols represent individual mice.

Similarly, adoptive transfer of female autoreactive T cells into female recipients failed to induce granulomas (Figures 6D-F). To distinguish T cell-intrinsic from host-dependent effects, male T cells were transferred into male or female recipients (Figure 6G). Female recipients exhibited greater adrenal CD4^+^ T-cell infiltration than males, with similar Th subset composition but higher absolute cell numbers (Figures 6H-I, Figure S4F-G). Despite this increased infiltration, female recipients failed to develop granulomas (Figures 6J-K). In contrast, both male and female EAA mice exhibited reduced corticosterone production in this model (Figure 6L). These findings demonstrate a sex-dependent uncoupling of adrenal pathology and function. Whereas females were largely resistant to EAA-induced granuloma formation, they were only partially protected from EAA-induced corticosterone insufficiency. Given the fact that the corticosterone production was higher in females than in males (Figure 6C, L), it is possible that low corticosterone levels support granuloma formation.

## Discussion

In this study, we established a murine model of AD that recapitulates a key hallmark of the human disorder – primary adrenal insufficiency. Although the EAA model exhibits a milder phenotype than human AD, it represents the first defined murine system to demonstrate progressive autoimmune-mediated adrenal dysfunction.

The principal strength of the EAA model is that it combines a defined self-antigen, pathogenic autoreactive CD4^+^ T-cell, and measurable adrenal dysfunction, thereby enabling mechanistic studies of disease initiation, progression, and therapeutic intervention. In contrast to spontaneous autoimmune models, the timing and magnitude of disease can be experimentally controlled, facilitating the dissection of immune pathways that regulate autoimmune adrenalitis. However, the EAA model does not phenocopy all aspects of human AD. Adrenal insufficiency remains incomplete in the EAA, with preservation of the mineralocorticoid-producing zona glomerulosa, allowing long-term survival without steroid replacement. Furthermore, human AD is likely more heterogeneous than EAA, with multiple genetic and environmental factors contributing to disease initiation and progression. Nevertheless, the EAA model captures key immunopathogenic processes underlying autoimmune adrenal destruction and provides a valuable platform for testing therapeutic strategies and investigating mechanisms that would be difficult to study directly in patients.

We identified CD4⁺ T cell-derived IFNG as a central driver of EAA pathogenesis. The key role of Th1 cells is supported by clinical data from AD patients, in which 21-OH-reactive CD4⁺ T cells isolated from peripheral blood have been shown to produce IFNG (19, 21). IFNG-driven T-cell responses are also characteristic of patients with APS-1 (35). Therapeutically, treatment of five APS-1 patients with ruxolitinib, an inhibitor of JAK1/2 kinases required for IFNG signaling, has been shown to reduce T cell-derived IFNG levels and improve multiple autoimmune manifestations (35). Because AD is a common feature of APS-1 (36, 37), these findings raise the possibility that inhibition of IFNG signaling could preserve adrenal steroidogenic function. Because the transition from subclinical adrenalitis to overt adrenal insufficiency is often slow and may be predicted by the presence of 21-OH autoantibodies (38, 39), targeted immunotherapy aimed at suppressing IFNG signaling may provide a window of opportunity to intervene early and prevent irreversible adrenal damage. While our data support a dominant Th1/IFNG axis, the presence of diverse CD4^+^ T-cell infiltrates suggests that additional effector subsets, such as Th17 cells, may also contribute.

Previous therapeutic efforts to restore endogenous glucocorticoid production in newly diagnosed AD patients, which combined B-cell depletion (rituximab) with ACTH-mediated adrenal stimulation, failed to reverse adrenal insufficiency (40). Our data from *Rag2*^-/-^ mice, which lack B cells and still develop robust disease, demonstrate that B cells are dispensable for EAA pathogenesis. These findings suggest that therapeutic strategies that suppress the effector functions of autoreactive CD4⁺ T cells, rather than B cells, have the potential to preserve adrenal function in AD.

IFNG-dependent granuloma formation in the EAA model, as well as granulomatous adrenal inflammation in tuberculosis-driven AD (32), underscores the critical role of Th1-mediated macrophage polarization in adrenal pathology. The functional significance of these granulomas, which accumulate progressively only in males and may occupy up to 20% of the adrenal cortex during EAA, remains unclear. The pro-inflammatory environment likely compromises adrenocortical cell viability (41), disrupts metabolic homeostasis (42), and impairs steroidogenic function. We speculate that excessive inflammation promotes adrenocortical cell destruction and accumulation of lipid-rich cellular debris that is inefficiently cleared by macrophages, leading to foamy macrophage aggregates. Consistent with this idea, lipid-laden foamy macrophages accumulate in the adrenal cortex of mice in an age- and diet-dependent manner (43).

In the two-step immunization EAA setting, females appear largely resistant to overt pathology, yet they exhibit measurable corticosterone insufficiency in the enhanced adoptive transfer model, albeit to a lesser extent than males and without granuloma formation. This indicates that while immune activation is sufficient to impair adrenal function in both sexes, the downstream tissue response diverges, with males showing a greater propensity for granulomatous remodeling. Because adrenal histopathology from AD patients is scarce, it remains unknown whether granuloma formation, although not a typical hallmark of AD, occurs in a subset of patients, including those with APS-1. Thus, the granulomatous phenotype observed in our model may represent a context-dependent consequence of CYP11A1-driven autoimmunity. Structurally similar syncytial lipoid aggregates have been observed in 18-week-old male mice harboring a *Cyp11a1* promoter mutation that causes adrenal-specific steroidogenic deficiency (44), suggesting that impaired steroidogenesis and chronic inflammation may converge on common tissue remodeling pathways. It is plausible that corticosterone inhibits granuloma formation, which would explain the absence of granulomas in female mice, which generally have higher corticosterone levels under steady-state and EAA conditions. Moreover, sex-related differences in macrophage composition (45) and the higher adrenocortical cell turnover observed in females (46) may further contribute to their relative resistance to granuloma formation.

To date, a robust and widely accepted murine model of AD has been lacking. Previous attempts, such as repeated immunization with adrenal extracts and lipopolysaccharide, elicited only mild cortical inflammation without inducing overt adrenal insufficiency, even after prolonged exposure (47). Consistent with this idea, adoptive transfer of CYP21-reactivated CD4⁺ T cells resulted in detectable infiltration of the adrenal glands without overt adrenal pathology. The inability of CYP21-reactive CD4⁺ T cells to induce adrenal insufficiency despite adrenal infiltration suggests that tissue infiltration alone is insufficient for disease development and that the antigen specificity determines the pathogenic potential of autoreactive CD4⁺ T cells. Autopsy studies reporting focal adrenal lymphocytic infiltrates in up to 7% of individuals younger than 54 years (48), far exceeding the clinical incidence of AD, further support the notion that many adrenal immune responses remain subclinical or abortive. Observed enhancement of corticosterone levels following peptide-free immunization may reflect transient systemic immune activation induced by the immunization regimen (poly(I:C) and anti-CD40), which could engage the hypothalamic-pituitary-adrenal axis and thereby increase adrenal steroidogenic responsiveness. Mitchell and Pearce proposed that high intra-adrenal glucocorticoid concentrations establish a locally immunosuppressive microenvironment, limiting immune effector function (1). Together with the presence of intraglandular regulatory T cells, this may explain the delayed onset of adrenal dysfunction in our model despite the accumulation of autoreactive CD4⁺ T cells.

Our findings identify CYP11A1 as a physiologically relevant adrenal autoantigen. Although clinical diagnosis relies on 21-OH autoantibodies, the prevalence of AD in APS-1 and the AIRE-dependent expression of *Cyp11a1* in mTECs (49) argue for its central role in tolerance and adrenal autoimmunity. ScRNA-seq revealed that *Cyp11a1* is expressed in a subset of AIRE⁺ mTECs, consistent with promiscuous gene expression observed in the thymus (50, 51), whereas *Cyp21a1* expression was more restricted. Although steroidogenic properties have been proposed for TECs (52), the enhanced susceptibility of *Aire*^-/-^ mice to EAA supports a critical role for central tolerance to CYP11A1. Three of the four tested adrenal-infiltrating T-cell clones recognized the CYP11A1_233-247_ self-peptide, thereby validating the bona fide autoimmune nature of EAA.

Our findings, together with evidence of progressive adrenal dysfunction and the intrinsic regenerative capacity of the adrenal cortex (46, 53, 54), position the adoptive EAA model as a valuable system for investigating the etiology of AD and provide an opportunity for preclinical testing of therapeutic interventions targeting autoimmune adrenalitis. The key role of IFNG-producing CD4⁺ T cells in EAA pathogenesis suggests that targeting IFNG signaling pathways may be a promising therapeutic strategy for AD in humans.

## Methods

### Sex as a biological variable

Our study examined male and female mice, and sex-dimorphic effects are reported.

### Mice

Male and female mice aged 6–13 weeks were used. Experimental groups were age- and sex-matched whenever possible, with littermates distributed evenly between groups. Mice were assigned to experimental groups based on animal ID before investigator involvement. *Cd3e*^-/-^ (RRID:IMSR_JAX:004177) (55), Ly5.1 (RRID: IMSR_JAX:002014) (56), *Aire*^-/-^ (RRID:IMSR_JAX:004743) (57), *Rag2*^-/-^ (RRID:IMSR_JAX:008449) (58), *Tnfr1*^-/-^(59) and *Ifng*^-/-^(60) strains were described. Mice were bred and maintained in a specific pathogen-free facility at the Institute of Molecular Genetics of the Czech Academy of Sciences (IMG). Mice were housed under a 12-h light/dark cycle with ad libitum access to food and water.

### Antibodies and Reagents

A complete list of antibodies and reagents used in this study is provided in the Supplementary Materials.

### Immunogenic Peptides

I-A^b^ MHCII-restricted CYP11A1 peptides (NCBI Reference Sequence: NP_031830.2) were designed using the Immune Epitope Database (IEDB.org). Two non-overlapping peptides with the highest predicted binding affinities were selected: CYP11A1_150-164_, VLNQEVMAPGAIKNF, and CYP11A1_233-247_, AVYQMFHTSVPMLNL. For CYP21, two candidate peptides (CYP21_143-158_, FCERMRAQPGTPVAI, CYP21_344-359_, VLRLRPVVPLALPHR) were evaluated by antigen recall assays using splenocytes from immunized C57BL/6 mice. Only CYP21_344-359_ elicited a robust response. An additional peptide, CYP21_372-386_, KDMVIIPNIQGANLD, was selected based on predicted I-Aᵇ binding affinity and used together with CYP21_344-359_ in adoptive transfer experiments. Peptides were synthesized at 70-85% crude purity with C-terminal amidation (Peptides & Elephants), dissolved in DMSO, aliquoted, and stored at -80°C as 20 mM stocks.

### Experimental Autoimmune Adrenalitis (EAA) Model

Mice aged 8-13 weeks were immunized subcutaneously at two dorsal sites with 100 µg of each peptide emulsified in CFA, prepared by supplementing Incomplete Freund’s Adjuvant (Sigma-Aldrich) with 4 mg/ml of heat-inactivated *Mycobacterium tuberculosis* H37RA (BD Difco). 10-14 days later, mice received an intraperitoneal booster containing 100 µg of each peptide, 50 µg high-molecular-weight poly(I:C) (InvivoGen), and 100 µg anti-CD40 antibody (clone FGK4.5). Control mice received the corresponding adjuvant formulations without peptide.

### Adoptive Transfer Model

B6 or *Ifng*^-/-^ mice were immunized with pCYP11A1 in CFA as described above. Ten days later, cells from the axillary, inguinal, cervical, and mesenteric lymph nodes were cultured for three days in complete RPMI (10% FBS, 100 U/ml penicillin (BB Pharma), 100 µg/ml streptomycin (CELLPURE)) containing pCYP11A1 (10 µg/ml). Recombinant IL-2 (2 ng/ml, Gibco) was added after 24 h. CD4⁺ T cells were enriched by magnetic separation (Miltenyi Biotec), stained for CD4, CD44, and CD62L, and CD4⁺CD44⁺ cells were sorted by flow cytometry. A total of 1 × 10⁵ cells were transferred intravenously into *Rag2*^-/-^ or *Cd3e*^-/-^ recipient mice. The following day, recipients underwent the standard EAA protocol. Control recipients received an equal number of CD4⁺CD44⁺ cells from CFA-immunized donor mice without peptide restimulation. Control donors were immunized three days later to synchronize cell harvest and transfer.

### Flow Cytometry and Cell Sorting

Adrenal glands were digested in complete RPMI containing collagenase D (0.5 mg/ml) and DNase I (0.1 mg/ml; Roche) at 37°C for 25 min with mechanical dissociation. Cell suspensions were filtered through a 70-µm nylon mesh, washed with HBSS containing 2% FBS and 2 mM EDTA, and stained for 30 min on ice with fluorochrome-conjugated antibodies, LIVE/DEAD Fixable Near-IR dye (Thermo Fisher Scientific), and anti-CD16/32 Fc block (clone 2.4G2). Intracellular staining was performed using the Foxp3/Transcription Factor Staining Buffer Set (Invitrogen). For intracellular cytokine staining, cells were stimulated with PMA (50 ng/ml) and ionomycin (1 µM) for 3-4 h in the presence of Golgi Block (Invitrogen). Where indicated, CD45.2-PE (clone 104) was administered intraperitoneally 15 min before euthanasia to distinguish tissue-resident from intravascular leukocytes. Samples were analyzed on a Cytek Aurora spectral cytometer, and cell sorting was performed using a FACSAria II or MoFlo Influx (BD Biosciences). Data were analyzed with FlowJo v10.8.0 (Tree Star).

### Adrenal Histology, Immunohistochemistry and Immunofluorescence

Adrenal glands were dissected free of surrounding fat under a stereomicroscope and fixed in 4% formaldehyde (Sigma-Aldrich) in PBS for 3 h. Samples were either processed for paraffin embedding or cryoprotected overnight in 30% sucrose (Sigma-Aldrich), embedded in OCT compound (Sakura), and stored at −80°C. Cryosections (10 µm) were prepared using a Leica CM1950 cryostat.

Paraffin sections (4 µm) were processed and stained with hematoxylin and eosin (H&E) at the Czech Centre of Phenogenomics using standard protocols and scanned with an Axio Scan.Z1 slide scanner (Zeiss). Where indicated, H&E staining was performed on cryosections using standard procedures and imaged on a Leica DM6000 widefield microscope.

For immunofluorescence, cryosections were fixed, permeabilized, and blocked before overnight incubation with fluorochrome-conjugated anti-CD45.2 (clone 104) and anti-CD4 (clone RM4-5) antibodies or rabbit anti-CYP11A1 antibody (clone D8F4F; 1:100), followed by Alexa Fluor 647-conjugated goat anti-rabbit secondary antibody (1:1000). Nuclei were counterstained with DAPI. Granuloma autofluorescence was detected using 488-nm laser excitation with emission detection 495-600 nm. Slides were mounted in ProLong Gold Antifade Mountant (Invitrogen) and imaged using a Leica TCS SP8 confocal microscope.

F4/80 immunohistochemistry was performed on formalin-fixed, paraffin-embedded adrenal sections at the Czech Centre of Phenogenomics using rabbit anti-F4/80 antibody (clone D2S9R; 1:750) with HRP/DAB detection according to standard protocols.

Neutral lipid accumulation was assessed by Oil Red O staining of frozen adrenal sections (8-10 µm) at the Czech Centre of Phenogenomics using standard procedures. Sections were counterstained with hematoxylin and mounted in Aquatex® aqueous mounting medium (Sigma-Aldrich).

### ACTH Stimulation Test, Serum Collection and Corticosterone Measurement

Synthetic ACTH (1-24: SYSMEHFRWGKPVGKKRRPVKVYP, 95% purity, Peptides & Elephants) was dissolved in distilled water (1 mg/ml). To suppress endogenous corticosterone production, mice received dexamethasone (0.1 mg/mouse, i.p.; Sigma-Aldrich) at 8:00 a.m. Two hours later, ACTH was administered intraperitoneally (1 mg/kg). One hour after ACTH injection, blood was collected from the superficial facial vein, allowed to clot for 30 min at room temperature, and centrifuged (1,250 × g, 15 min, 4°C). Serum was collected and stored at −80°C. Mice were allowed to recover for at least three days before subsequent analyses.

Serum corticosterone concentrations were measured using a commercial ELISA kit (Tecan) according to the manufacturer’s instructions. Absorbance was measured using an EnVision plate reader (PerkinElmer), and data were analyzed using MyAssays software.

### Na^+^/K^+^ measurement

Serum sodium and potassium concentrations were measured using an AU480 automated biochemical analyzer (Beckman Coulter) at the Czech Centre of Phenogenomics according to the manufacturer’s protocols.

### scRNA-seq

B6 mice (10-12 weeks old) were immunized with pCYP11A1 in CFA using the protocol described above, whereas control mice received CFA alone. Fourteen days later, adrenal glands and spleens were harvested. Adrenal glands were processed as described for flow cytometry, whereas splenocytes were processed without enzymatic digestion. Prior to cell sorting, adrenal CD4⁺ T-cell infiltration was quantified by flow cytometry using anti-TCRβ (clone H57-597) and anti-CD4 (clone H129.19) antibodies. Samples with the highest adrenal infiltration were selected for scRNA-seq. Cells were stained with anti-TCRβ, anti-CD4, anti-CD11b (clone M1/70), and TotalSeq-C Hashtag antibodies (BioLegend) for sample multiplexing. Live CD11b⁻TCRβ⁺CD4⁺ T cells were sorted using a MoFlo Influx cell sorter. Cell viability exceeded 90% before loading onto the Chromium Controller (10x Genomics).

Single-cell 5′ gene expression, VDJ, and Feature Barcode libraries were prepared using Chromium Single Cell 5′ reagents (10x Genomics) according to the manufacturer’s instructions and sequenced on an Illumina NovaSeq 6000.

Sequencing data were processed using Cell Ranger v5.0.1 and aligned to the *Mus musculus* reference genome (GRCm38, Ensembl release 102) (61). Downstream analyses were performed in R v4.2.1 using Seurat v4.3.0 (62). Cells lacking unique hashtag assignment, cells with multiple hashtags, and cells with high mitochondrial content or identified as doublets were excluded. Adrenal and spleen datasets were analyzed separately, integrated using Seurat’s canonical correlation analysis (CCA), clustered, and manually annotated. The final dataset comprised 5,584 CD4⁺ T cells.

Gene set enrichment analysis was performed using fgsea with Hallmark gene sets from the Molecular Signatures Database (MSigDB). Heatmaps were generated using pheatmap.

TCR repertoires were reconstructed using Cell Ranger VDJ and MiXCR v3.0.13 (62) with the murine IMGT reference (63). Cells with multiple productive TCRβ chains or excessive TCRα/TCRβ rearrangements were excluded as doublets. Cells sharing identical paired TCRα and TCRβ sequences were classified as clones.

### Transduction of A5 cells with ADR-specific TCRs

A5 hybridoma cells were kindly provided by Ludger Klein (Institute for Immunology, Biomedical Center (BMC), Faculty of Medicine, LMU Munich, Planegg-Martinsried, Germany). Cells were maintained in complete IMDM supplemented with 5% FBS, penicillin (100 U/ml), and streptomycin (100 µg/ml). Cells (4 × 10⁶) were transduced with retroviral supernatants encoding pMSCV-ADR1-4-LNGFR or pMSCV-B3K508-LNGFR in the presence of polybrene (10 µg/ml) by spinoculation (2500 rpm, 45 min). After overnight incubation at 37 °C, cells were transferred to fresh medium. At 48 h post-transduction, LNGFR-expressing cells were stained with CD271 (LNGFR)-AF647 antibody (clone ME20.4) and sorted by FACSAria II (BD Biosciences). Sorted cells (>90% purity) were cryopreserved in FBS containing 10% DMSO.

### Monoclonal Retrogenic T cells

Generation of immortalized bone marrow progenitors was performed as previously described (64, 65). Briefly, *Rag2*^-/-^ mice were treated with 5-fluorouracil (100 mg/kg), and bone marrow cells were harvested 5 days later. Cells were cultured with stem cell factor, IL-3, and IL-6 and transduced with a NUP98-HOXB4 fusion construct in the pMYs retroviral vector. Transduced progenitors were selected with puromycin and used for TCR transduction.

For generation of monoclonal CD4⁺ T cells, immortalized progenitors were transduced with GFP-expressing MSCV vectors encoding TCRα and TCRβ chains derived from ADR1-ADR4 clones identified by scRNA-seq. GFP⁺ cells were sorted by flow cytometry and transplanted intravenously into lethally irradiated congenic Ly5.1 recipients. After 8 weeks, GFP⁺Ly5.2⁺CD4⁺ T cells were isolated by FACS and used for antigen presentation assay.

### Antigen presentation assays

Monoclonal CD4⁺ T cells isolated from retrogenic mice were co-cultured with Ly5.1 splenocytes as antigen-presenting cells (APCs) at a 3:1 responder-to-APC ratio in complete RPMI medium. Cells were stimulated for 24 h with individual pCYP11A1 peptides (10 µM) or left unstimulated, followed by flow cytometric analysis of activation markers. Dead cells were excluded using LIVE/DEAD Fixable Near-IR viability dye.

For antigen-specific activation, ADR1-4 A5 cells were co-cultured with B6 splenocytes (1 × 10⁵ cells each) in 96-well U-bottom plates and stimulated with pCYP11A1 across a titrated concentration range (10 µM-10 pM) for 24 h. Cells were then stained with LIVE/DEAD NIR, CD4-AF594 (clone GK1.5), and CD271-AF647 (clone ME20.4) prior to analysis. Controls included transduced cells cultured alone, co-culture without peptide, and irrelevant peptide stimulation. PMA and ionomycin treatment (4 h) served as a positive control for TCR-independent activation.

### In vitro recall assays

To assess peripheral CYP21- versus CYP11A1-specific T cell responses, splenocytes from previously immunized B6 mice were cultured in complete RPMI medium and restimulated with peptide pools containing pCYP11A1 or pCYP21, or with individual CYP21-derived peptides (10 µM each), for 24-48 h. Supernatants were collected, and IFNG levels were quantified by ELISA (BioLegend) according to the manufacturer’s instructions.

### Data and code availability

The raw sequencing data have been deposited in the NCBI Gene Expression Omnibus (GEO) under accession number GSE303645. The code used for analysis has been deposited on GitHub (https://github.com/Lab-of-Adaptive-Immunity/Project_Adrenalitis). Values for all data points in the graphs are provided in the Supporting Data Values file.

### Statistical analyses

Data were analyzed by two-tailed nonparametric Mann-Whitney test with GraphPad5 software. Data are presented as mean ± SEM. Each symbol represents an individual mouse. P value less than 0.05 was considered significant.

### Study approval

All animal experiments were approved by the Expert Committee for Animal Welfare of the Czech Academy of Sciences, Prague, Czech Republic (approval nos. AVCR 6917/2022 SOV II and AVCR 4025/2026 PPZ). All procedures were conducted in accordance with the ARRIVE Essential 10 guidelines.

## Acknowledgement

We gratefully acknowledge Ladislav Cupak for technical assistance, Zdenek Cimburek and Matyas Sima (Flow cytometry facility, IMG) for cell sorting, Sarka Kocourkova and Michal Kolar (Laboratory of Genomics and Bioinformatics, IMG) for cDNA library preparations, Jachym Antonin Harwood (IMG) for advice on the manuscript, and Dominik Filipp (IMG) for providing us with *Aire*^-/-^ mice. AA, VC, and VN are students of the Faculty of Science Charles University in Prague. We acknowledge the Light Microscopy Core Facility, IMG, Prague, Czech Republic, supported by MEYS – LM2023050, MEYS – CZ.02.1.01/0.0/0.0/18_046/0016045 and MEYS – CZ.02.01.01/00/23_015/0008205, for their support with the confocal/widefield/imaging and image analysis presented herein. Illustrations were created using Biorender.

## Author contributions

**Arina Andreyeva:** Formal analysis; Investigation; Visualization; Methodology; Writing. **Juraj Michalik:** Data curation; Software; Formal analysis; Writing. **Veronika Niederlova:** Investigation; Visualization. **Veronika Cimermanova:** Investigation; Visualization. **Ales Drobek:** Investigation, Methodology. **Radislav Sedlacek:** Resources; Supervision; Methodology**. Jan Prochazka:** Resources; Supervision; Methodology **Juraj Labaj:** Resources; Supervision; Methodology. **Olha Fedosieieva:** Investigation, Methodology. **Waldemar Kanczkowski:** Investigation; Methodology. **Peter Draber:** Resources. **Andre Sulen**: Investigation; **Ondrej Stepanek:** Conceptualization; Visualization; Resources; Supervision; Funding acquisition; Writing. **Ales Neuwirth:** Conceptualization; Supervision; Formal analysis; Investigation; Visualization; Methodology; Writing.

## Declaration of Interests

The authors declare no competing interests.

## Funding support

- EU Horizon 2020 research and innovation programme, ERC Starting Grant ActSwiftly, No. 101125695 (to OS).
- Czech Science Foundation, No. 22-18046S (to OS)
- Charles University Grant Agency, No.176624 (to VC)
- Swiss National Science Foundation, No. 320030-227564 (to PD)
- Institute of Molecular Genetics of the Czech Academy of Sciences, institutional support No. RVO 68378050
- Czech Centre for Phenogenomics, infrastructure support No. LM202303 (Ministry of Education, Youth and Sports of the Czech Republic).
- Operational Programme Research, Development and Education (OP RDE), No. CZ.02.1.01/0.0/0.0/16_013/0001789, Upgrade of the Czech Centre for Phenogenomics: Developing Towards Translational Research (Ministry of Education, Youth and Sports of the Czech Republic and the European Structural and Investment Funds).
- Light Microscopy Core Facility, Institute of Molecular Genetics of the Czech Academy of Sciences, supported by the Ministry of Education, Youth and Sports of the Czech Republic through projects No. LM2023050, No. CZ.02.1.01/0.0/0.0/18_046/0016045, and No. CZ.02.01.01/00/23_015/0008205.

## Notes

### Competing Interest Statement

The authors have declared no competing interest.

### Summary of Updates

This revision included novel experiments and changes in the text made during a revision process for a peer-reviewed journal. One new author and several funding sources were added.

