## Supplementary material for "IFNG-producing self-reactive CD4^+^ T cells induce autoimmune adrenalitis in a mouse model of Addison’s disease": Figure S

Supplementary figure 1

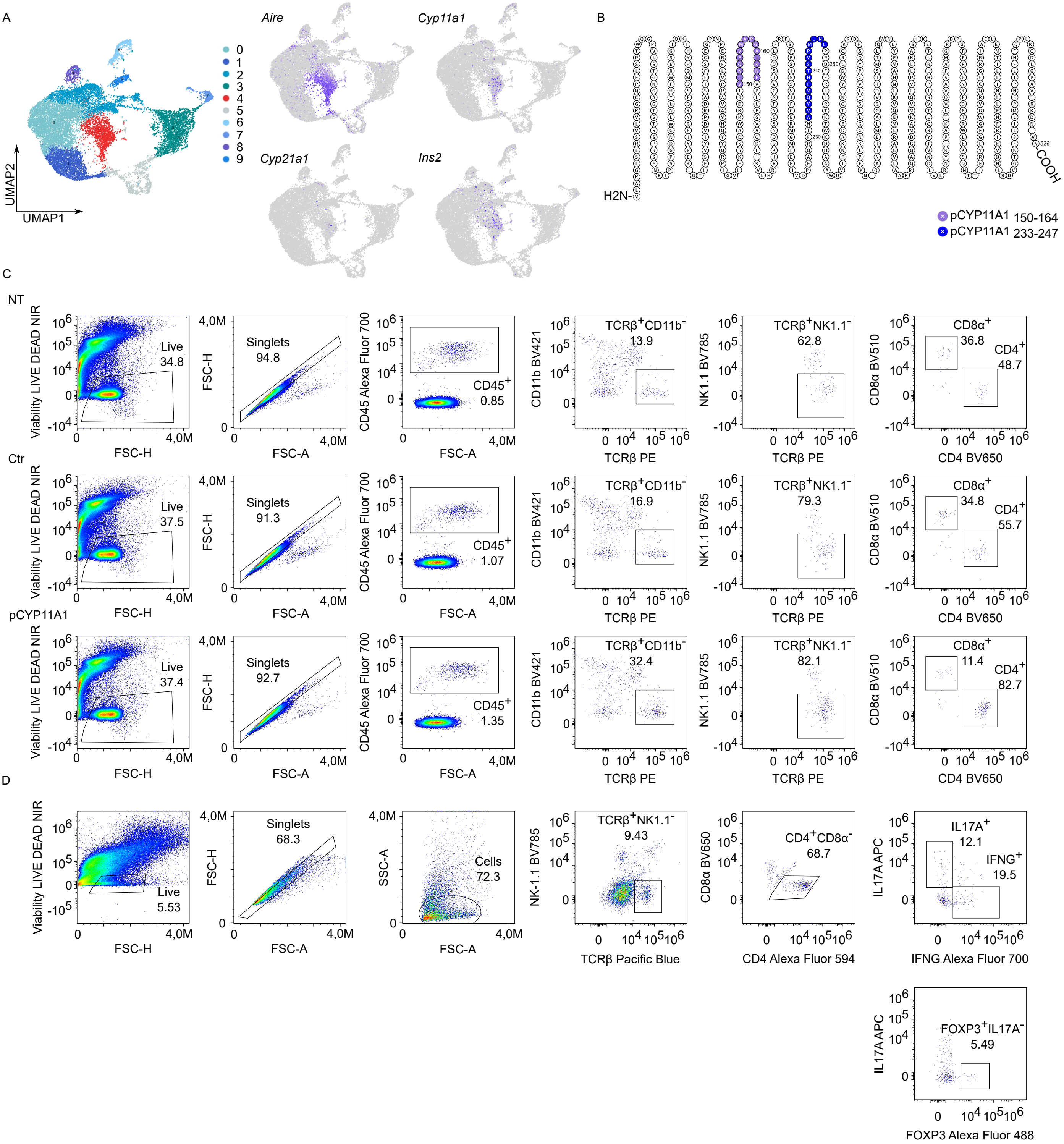

**Supplementary Figure 1. *Cyp11a1* expression in mTECs, immunogenic CYP11A1 peptides and identification of adrenal CD4$^+$ T cells.**

**(A)** UMAP of scRNA-seq data from TECs of 8-week-old female mice (also in Figure 1A) with feature plots displaying expressions of selected genes and highlighting Aire-expressing mTECs (cluster 4). Gene expression is visualized in shades of violet, with higher intensity reflecting greater expression levels.

**(B)** Visualization of the linear protein sequence of murine CYP11A1 (Protter software) (Omasits et al., 2014) with the two peptides used for immunization - CYP11A1$_{150\text{-}164}$ and CYP11A1$_{233\text{-}247}$ - highlighted in indicated colors. Peptides were selected based on predicted immunogenicity and MHC class II binding potential.

**(C)** Representative gating for identifying adrenal CD4$^+$ and CD8$^+$ T cells (Figure 1B).

**(D)** Representative gating for identifying Tregs and cytokine-producing CD4$^+$ T cells (Figure 1C).

Supplementary figure 2

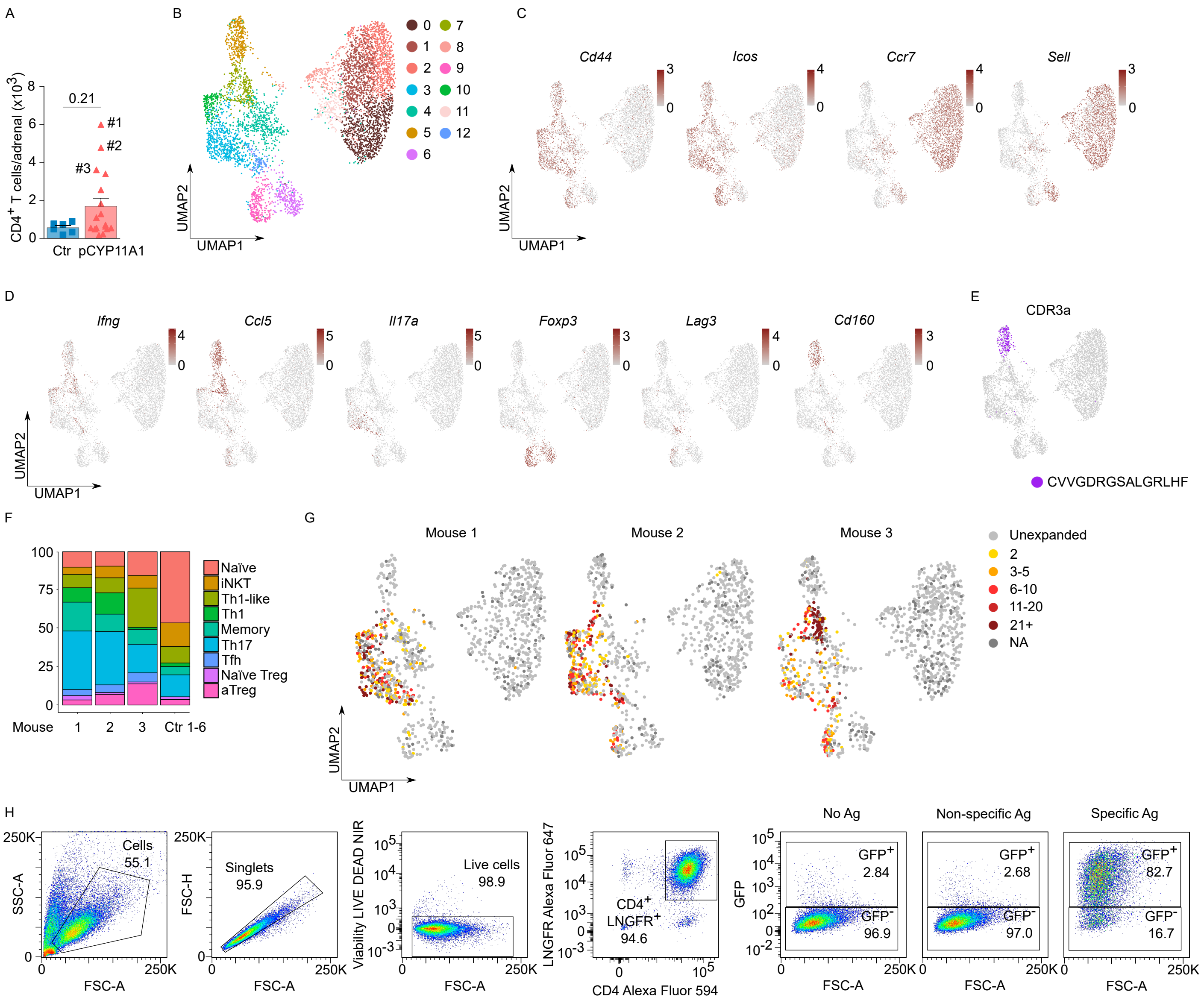

**Supplementary Figure 2. Heterogeneity and antigen specificity of adrenal CD4$^{+}$ T cells**

**(A**) Quantification of CD4$^{+}$ T-cell infiltration in adrenal glands of pCYP11A1-immunized and control mice at day 14 p.i. (n=6-17 mice/group).

Statistical significance was determined using two-tailed Mann-Whitney test; P values are indicated. Bars show the mean ± SEM; individual mice are shown as symbols.

**(B)** UMAP of scRNA-seq data showing 13 transcriptionally defined CD4$^{+}$ T cell clusters from adrenal glands and spleens. Five similar clusters with a naïve phenotype were merged and labeled in the final projection as "Naïve CD4$^{+}$" (Figure 1D).

**(C)** Expression of signature genes for naïve and effector CD4$^{+}$ T cell subsets in cells derived from adrenal glands and spleens.

**(D)** Expression of marker genes used to identify specific effector CD4$^{+}$ T cell subsets in cells derived from adrenal glands and spleens. In (C-D) color intensity correlates with the level of gene expression as indicated.

**(E)** Feature plot identifying the CD4$^{+}$ invariant NKT (iNKT) cell subset based on expression of the canonical CDR3α sequence (CVVGDRGSALGRLHF).

**(F)** Distribution of transcriptionally defined CD4$^{+}$ T-cell populations in adrenal glands only, shown for individual immunized mice (n = 3 mice) and pooled control mice (n = 6 mice).

**(G)** UMAP showing clonal expansion of CD4$^{+}$ T cells in adrenals and spleen from individual pCYP11A1-immunized mice. Color represents clone size, defined by the number of cells sharing the same TCR; "NA" denotes T cells without complete information about both TCR chains.

**(H)** Representative gating strategy for NFAT-GFP reporter T cell hybridoma (A5) clones used for antigen specificity assays (Figure 2G).

Supplementary figure 3

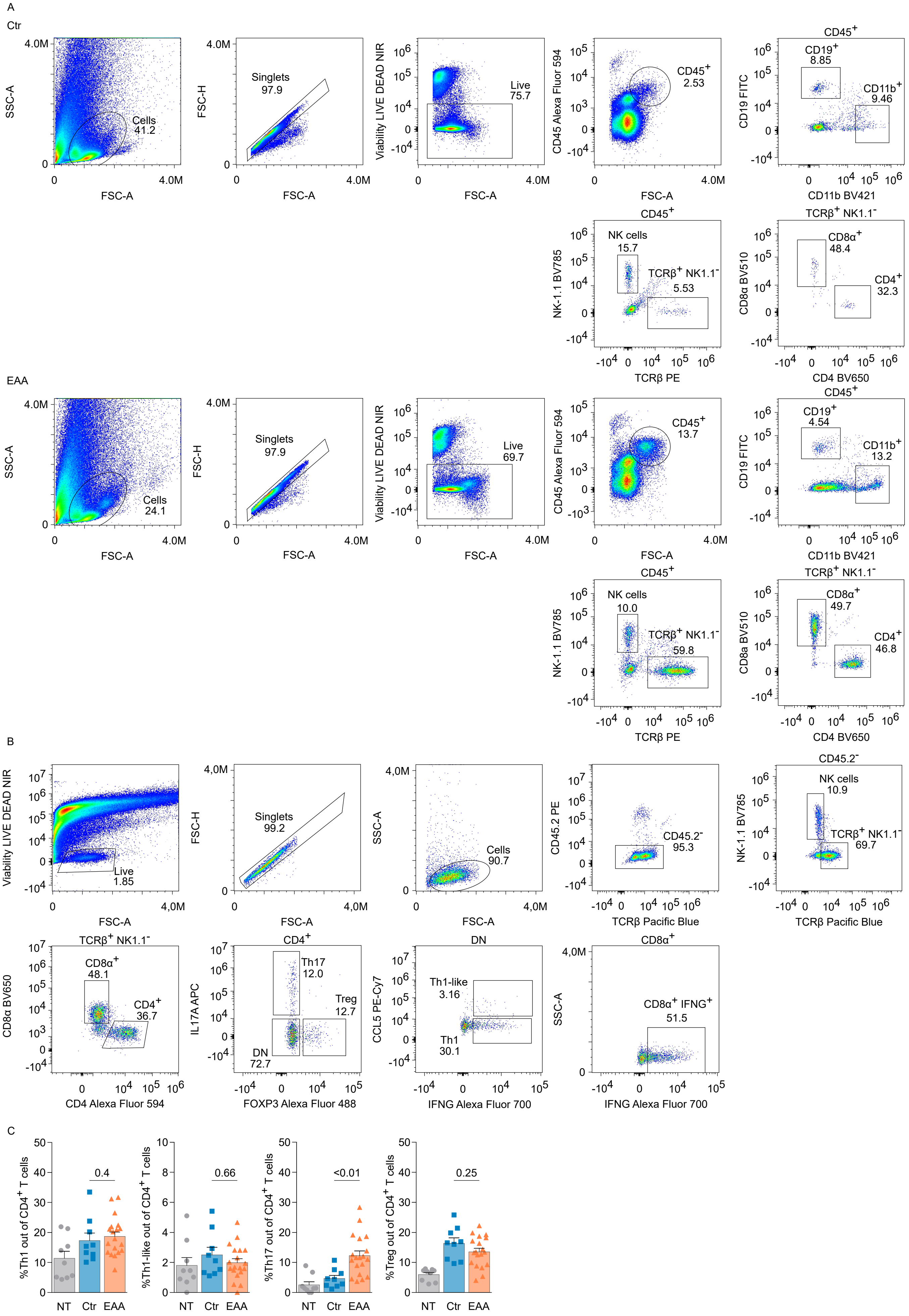

**Supplementary Figure 3. Immune cell characterization in the adrenal glands in the EAA model.**
**(A)** Representative gating for identifying major immune cell populations in adrenal glands of Ctr and EAA mice (Figure 3A).
**(B)** Representative gating for identifying $CD4^+$ effector T cell subsets (Figure 3C-D). CD45.2 PE antibody was injected 15 minutes prior to sacrifice, and blood-derived cells were excluded by gating on $CD45.2^-$ cells.
**(C)** Frequencies of $CD4^+$ effector T-cell subsets 3-4 weeks p.i., shown as percentages of total $CD4^+$ T cells (two independent experiments; n=9-19 mice/group).
Groups: non-treated (NT), control (Ctr), and EAA.
Statistical significance was determined using two-tailed Mann-Whitney test; P values are indicated. Bars show the mean ± SEM; individual mice are shown as symbols.

Supplementary figure 4

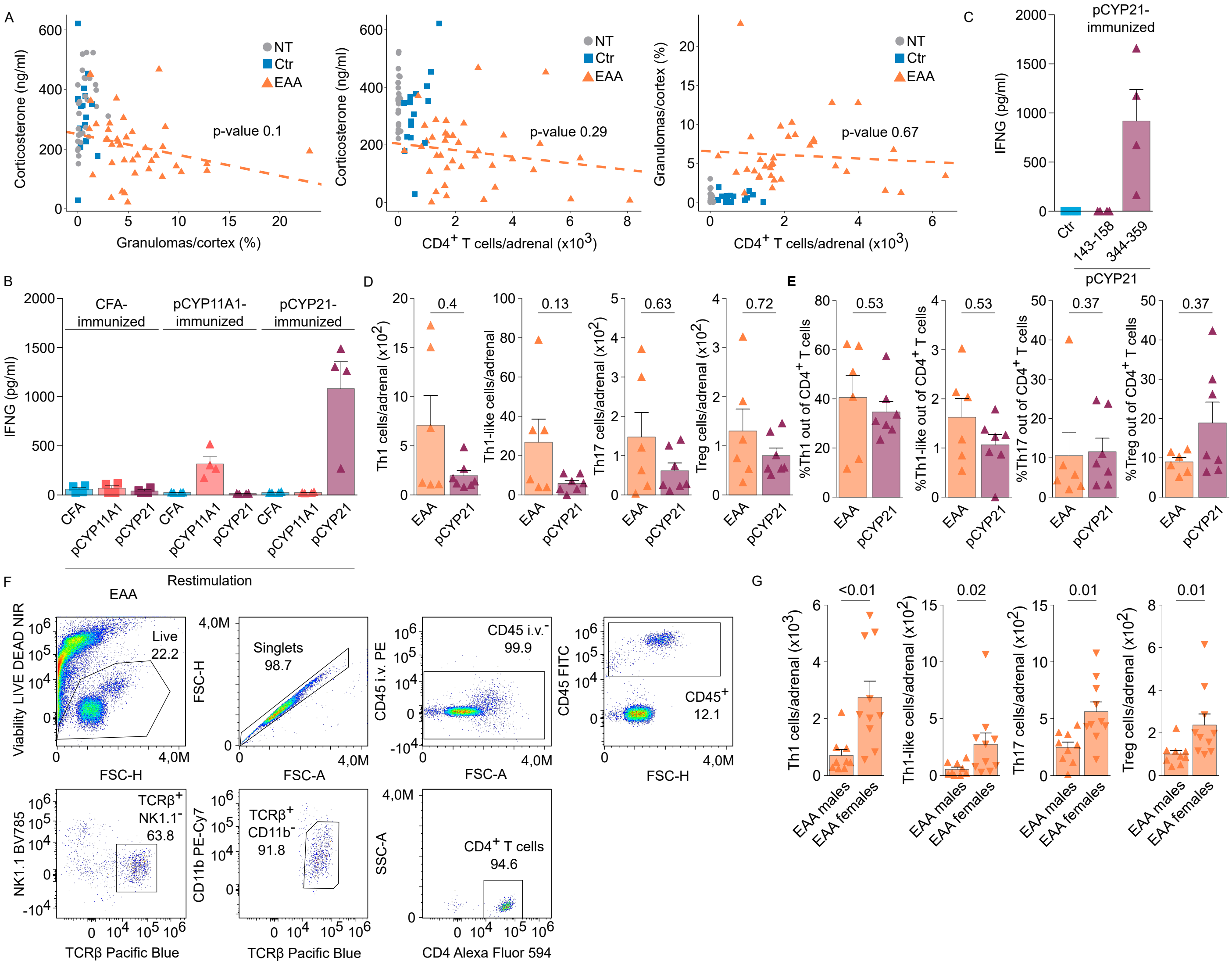

**Supplementary Figure 4. Antigen-specific and sex-dependent features of adrenal pathology in the EAA model.**

**(A)** Correlation between serum corticosterone levels, granuloma frequency, and $CD4^+$ T-cell infiltration in the adrenals of recipient mice (*Cd3e*$^{-/-}$, *Rag2*$^{-/-}$) in the adoptive EAA model 3 months p.i.

**(B)** IFNG levels in supernatants of splenocytes after restimulation with pCYP11A1 or pCYP21 ($CYP21_{143-158}$, $CYP21_{344-359}$) from previously pCYP11A1-or pCYP21-immunized B6 mice.

**(C)** IFNG levels in supernatants of splenocytes after restimulation with individual CYP21 peptides ($CYP21_{143-158}$, $CYP21_{344-359}$).

**(D)** Absolute numbers and **(E)** frequencies of $CD4^+$ T-cell effector subsets infiltrating adrenal glands of *Cd3e*$^{-/-}$ recipients 3 months post-transfer of pCYP11A1 (EAA)-or pCYP21-restimulated $CD4^+$ T cells (two independent experiments; n=6-7 mice/group)

**(F)** Representative gating strategy for identification of $CD4^+$ T cells (Figure 6H). CD45.2 PE antibody was injected 15 minutes prior to sacrifice, and blood-derived cells were excluded by gating on $CD45.2^-$ cells.

**(G)** Absolute numbers of $CD4^+$ T-cell effector subsets infiltrating adrenal glands of *Rag2*$^{-/-}$ male and female recipients 3 months post-transfer (two independent experiments; n=9-10 mice/group). Cells were gated as in Figure S3D.

Omasits, U., C.H. Ahrens, S. Muller, and B. Wollscheid. 2014. Protter: interactive protein feature visualization and integration with experimental proteomic data. *Bioinformatics* 30:884-886.
